# Dynamical differential covariance recovers directional network structure in multiscale neural systems

**DOI:** 10.1101/2021.06.18.448901

**Authors:** Yusi Chen, Burke Q. Rosen, Terrence J. Sejnowski

## Abstract

Investigating neural interactions is essential to understanding the neural basis of behavior. Many statistical methods have been used for analyzing neural activity, but estimating the direction of network interactions correctly and efficiently remains a difficult problem (1). Here, we derive dynamical differential covariance (DDC), a new method based on dynamical network models that detects directional interactions with low bias and high noise tolerance under nonstationarity conditions. The method was first validated and compared with other methods on networks with false positive motifs and multiscale neural simulations where the ground truth connectivity was known. When applied to recordings of resting-state functional magnetic resonance imaging (rs-fMRI) recordings, DDC consistently detected regional interactions with strong structural connectivity in over 1,000 individual subjects obtained by diffusion MRI (dMRI). DDC is a promising new family of methods for estimating functional connectivity that can be generalized to a wide range of dynamical models and recording techniques and to other applications where system identification from sparse data is needed.

**Significance Statement:** Through dynamical interactions with each other, neurons make it possible for us to sense, move and think. It is now possible to simultaneously record from many individual neurons and brain regions. Methods for analyzing these large-scale recordings are needed that can reveal how the patterns of activity give rise to behavior. We developed an efficient, intuitive and robust way to analyze these recordings and validated it on simulations of model neural networks where the ground truth was known. We called this method dynamical differential covariance (DDC) because it can estimate not only the presence of a connection but also which direction the information is flowing in a network between neurons or cortical areas. We also successfully applied DDC to brain imaging data from functional Magnetic Resonance Imaging.

## 1. Introduction

Understanding the neural mechanisms underlying behavior requires knowing how neurons interact with each other. Direct circuit tracing by connectomics studies (2–4) provides ground truth, but at high cost and the resulting static connection matrix does not by itself reveal the dynamical aspects of neural communication. This has motivated statistical methods (1, 5) to estimate functional connectivity (FC) from recordings of neurons and brain imaging. (We refer here to all methods based on correlation or causation as FC (6)).

FC is often evaluated by estimating pairwise covariance, a symmetric measure that cannot detect directional coupling or disambiguate two unconnected nodes confounded with high correlation due to a common input (1, 7). Covariance estimates can be improved by using a suitably regularized partial covariance matrix, which is a global method that can “explain away” some of the ambiguities, but this matrix is also symmetric and does not solve the problem of finding the direction of coupling. Another assumption that covariance methods make is that conditions are stationary so that sample covariance can be obtained averaging across time, but this precludes changes in the level of brain arousal or tasks when switches occur between brain states. Despite these limitations, covariance methods have served as intuitive measures of brain coordination and remain the dominant way that most researchers estimate FC.

Another class of methods for estimating FC is based on the degree to which one time series can predict another time series. These methods include variants of Granger causality (8), cross convergent mapping (CCM) (9) and cross-dynamical delay differential analysis (cd-DDA) (10). For example, for Granger causality, time series A is causally related to time series B if the removal of A reduces the accuracy of predicting the future of B. As evident from the definition, selecting a best-fit model for prediction is essential. As a consequence, this class of methods can in principle improve FC estimates, but they require much more computation than covariance methods and do not scale well.

Other generative methods for FC such as dynamic causal modeling (DCM) (11) and Bayes net models (12, 13) search all possible causal graphs and fit the entire dataset to every hypothesis. These methods make further statistical assumptions and require even more computation, severely limiting the size of the network that can be analyzed.

Despite these advances in the statistical analysis of time series (6), we do not yet have an accurate and intuitive way to compute directional connectivity from brain recordings that is computationally efficient and statistically robust. One of the reasons for this shortcoming is that current methods are not derived from the processes that generate network activities.

We previously introduced differential covariance (dCov) (14, 15) (16), a directed FC estimation method, and highlighted the performance of two matrices, Δc, which is the correlation between the derivative signal and the signal itself, and Δp, which is the partial covariance between them. In simulated test cases, dCov detected network connections with higher sensitivity than many of the methods reviewed in Smith et al (1). In this paper, we derive a direct link between dCov and dynamical models of network activity. This leads to a new class of estimators called dynamical differential covariance (DDC) that explicitly defines FC as the interaction matrix that appears in the equations for a dynamical system.

In the following sections, we first analytically show that DDC provides unbiased estimates regardless of the noise structure, without assuming that the data are stationary. These favorable statistical properties were numerically confirmed in networks with both linear and nonlinear dynamics. We then show that DDC can infer the ground truth connections and their direction in multiscale neural network simulations. Lastly, we apply DDC to resting-state fMRI recordings from over 1,000 subjects and show that the extracted FC closely matches the structural connectivity measured by diffusion MRI.

## 2. Results

### A. Dynamical differential covariance (DDC)

Models of neural systems span a wide range of scales. At the microscopic level, the voltage trace, calcium dynamics and firing rate of a single neuron are highly nonlinear. These dynamics are often modeled using biophysical models based on voltage-gated ion channels. In contrast, at the macroscopic level the collective activity of a population of neurons and interactions between brain regions can be approximated by linear dynamics because of ensemble averaging (5, 17–19). For example, Nozari et al (19) showed that compared to other sophisticated non-linear families of models, the simple linear auto-regressive model performed best on modeling functional magnetic resonance imaging (fMRI) and intracranial electroencephalography (iEEG) recordings from hundreds of human subjects.

We first propose a linear dynamical model in Eq. 1 for global recordings and a nonlinear dynamical model in Eq. 2 for local neural recordings:

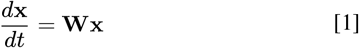

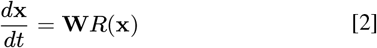

The column vector **x** is the neural activity, such as the membrane voltage or fMRI signal, **W** is the square connectivity matrix and *R*(**x**) is a nonlinear response function. Combining the above equations with **x** yields:

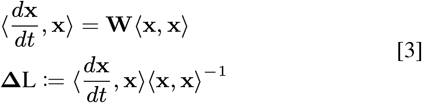

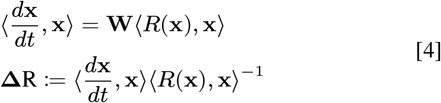

where **Δ**L and **Δ**R are DDC estimators for **W** and the operator ⟨, ⟩ takes the outer product of two vectors and performs time averaging.

The origin of the linear DDC estimator **Δ**L from a dynamical model provides an intuition for its effectiveness in estimating **W** as the product of two matrices: The first is differential covariance, which carries information about sources and sinks. In a neuron the sink is the inward current from synaptic inputs in the receiving area and in brain imaging it is related to changes in surrogates for local brain activity, whereas the source is the activity level in the sending area; In the second term, an entry in the partial covariance matrix is zero if and only if the residual correlation between **x**_**i**_ and **x**_**j**_ is zero, which cancels the influence of common sources. Robust estimation of directional interactions becomes possible by combining signals from sources and sinks and canceling signals from common sources. The multiplicative combination of the two terms yields better estimates than either alone.

A family of estimators arises from the DDC estimator **Δ**R for nonlinear dynamical systems for different *R*(**x**). Estimators can be adapted to the filtering effects from different recording techniques, such as the slow kinetics of calcium signals, by choosing the nonlinear function *R*(**x**) appropriately. Here, we use the rectified linear unit (ReLU), parameterized by a threshold, *θ*, that is often used in artificial neural networks (20), yielding ΔReLU as the corresponding nonlinear DDC estimator. The threshold can be set to optimize performance. Intuitively, the ReLU function rectifies low magnitude “noise” and retains larger signals.

### B. DDC provides unbiased estimation for nonstationary data

DDC estimators have several favorable statistical properties. First, given the correct neural model for the generative process, DDC provides an unbiased estimation of the connectivity matrix. In the Material and Methods G.1, we show analytically that in systems governed by stochastic differential equations, DDC gives unbiased estimates of connectivity **W** in the sense that the DDC estimation converges to the ground truth given sufficient number of trials. In addition, the estimation remains unbiased regardless of the noise structure (**D** in Eq. 10) - whether or not the added noise is correlated. To numerically confirm this result, we simulated a confounder network governed by linear dynamics (Fig. 1A) and estimated the connectivity from simulated time traces (Fig. 1B). At first glance, ΔL accurately recovered the ground truth network structure. To further investigate the origin of error, we orthogonally decomposed the total error (Methods) across trials into bias and variance (Fig. 1D). The bias part is due to the intrinsic property of the estimation method while the variance part drops as the number of estimation trials increases. Across 50 simulations, ΔL achieved the smallest estimation error, and more importantly, its bias over variance ratio (*θ*_*b*_) was also the lowest (Fig. 1D). Thus, both analytical and numerical results confirmed that DDC estimation is unbiased.

**Fig. 1.**
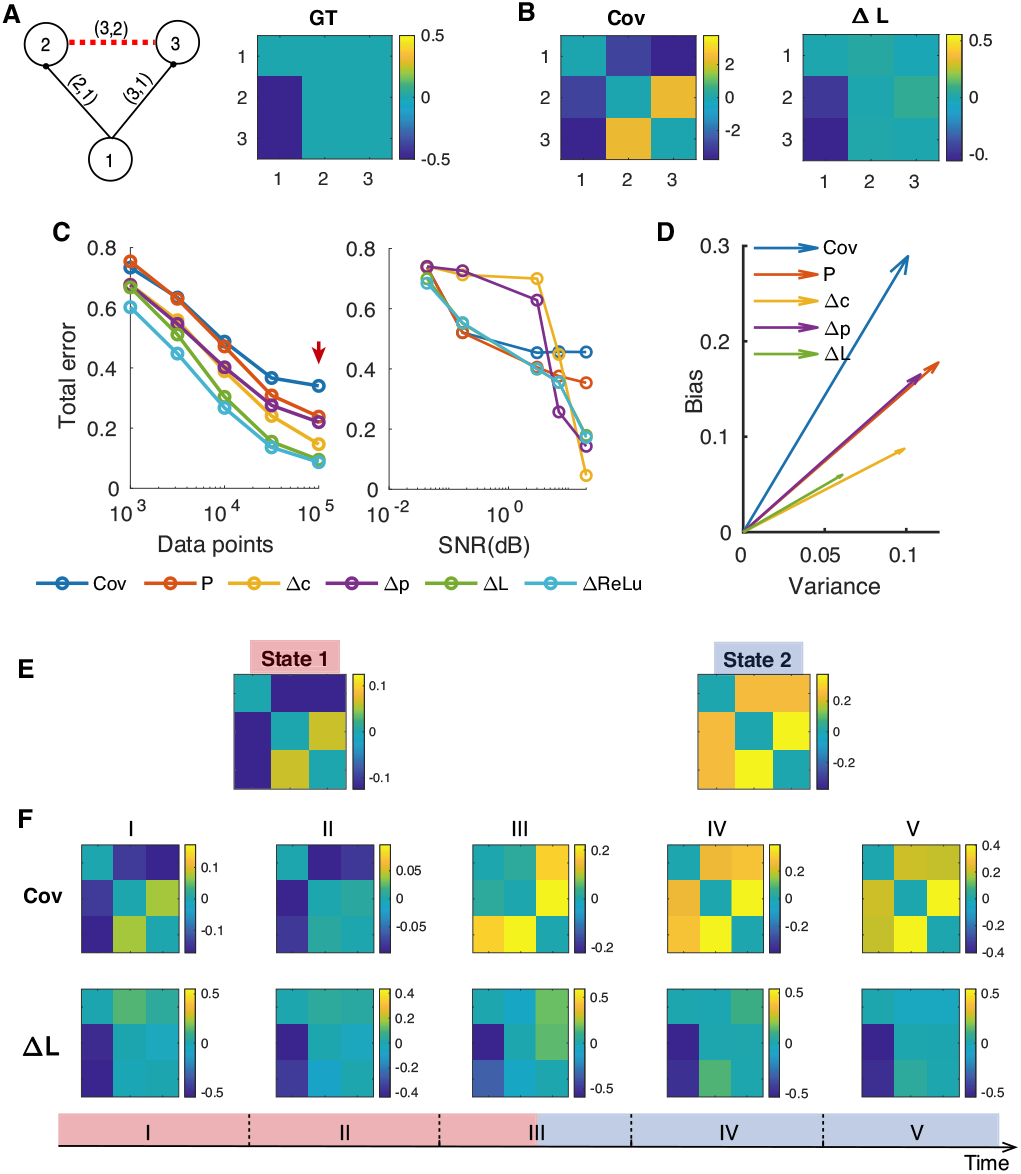
DDC provides unbiased estimation for nonstationary data. A) Left: Ground truth (GT) network structure. Black solid lines are directed physical connections and red dashed lines are false positive connections commonly inferred by covariance estimation; Right: GT shown in matrix form. B) Sample covariance (Cov) and ΔL estimation from time traces simulated by linear dynamics. C) Left: Influence of data size. Right: Influence of observational noise. SNR: signal to noise ratio; P: partial covariance estimation; Δc: differential covariance matrix; Δp: partial differential covariance matrix. D) Decomposition of estimation errors computed using sufficient data from 50 trials (corresponding to the red arrow in C). (E)(F) Simulation of the static confounder motif governed by a two-state dynamical system. E) Analytical solution of the steady-state covariance matrix in State 1 (left) and State 2 (right). F) Bottom: Timeline for sample time series with five non-overlapping sampling windows. The simulated time series was in State 1 during windows I and II and in State 2 during windows IV and V. Window III included data from both states. Sample covariance estimation (first row) shifted between states; in contrast, ΔL estimation (second row) consistently reported the static true connectivity. For a clear illustration, we removed the diagonal values from estimated matrices.

We also quantified the variance and bias over a range of data size and observational noise. DDC consistently had the least estimation error regardless of the size of the dataset (Fig. 1C). In contrast, inference bias for Covariance (Cov), Partial covariance (P), differential covariance (Δc) and partial differential covariance (Δp) diverged as data volume increased, introducing a systematic error. Regarding noise tolerance (Fig. 1C), the performance of dCov matrices (Δc and Δp) rapidly deteriorated with increasing noise probably due to the computation of derivative. However DDC remained robust.

Secondly, we also prove that DDC can be used to analyze nonstationary data whose higher-order statistics vary with time. In practical neural data processing, stationarity is often assumed when estimating the covariance matrix through sampling over time. However, this may not be a valid assumption because neural recordings can be quite nonstationary due to fluctuating brains states owing to neuromodulation and varying sensory inputs. There is need for methods that can analyze nonstationary data. In the derivation of DDC, stationarity was not required because relationship imposed by the system equation holds at every time step, regardless of the probability distribution of the process. To verify that DDC does not depend on stationarity, we simulated a two-state dynamical system (Methods) whose connectivity matrix (shown in Fig. 1A) remained time invariant while the noise structure was switched between states. The analytical solution of the covariance structure is given by Eq. 18, as plotted in Fig. 1E. Using non-overlapping sliding windows, we obtained the sample covariance and ΔL estimation of the connectivity matrix (Fig. 1F). The time-varying covariance matrix showed that the time trace is not stationary and thus sample covariance estimation fails to capture the true connectivity profile. On the other hand, ΔL consistently and accurately estimated the true connectivity matrix, even in window III where state switching occurred.

### C. DDC inferred the existence and direction of multiple network structures and dynamics

As a proof of principle, we applied DDC to three-node networks with varying dynamics and network structures (Fig. 2). The chain motif (Fig. 2A,B) and confounder motif (Fig. 2C) were chosen because they both have a node pair (red dashed line) that is highly correlated but with no physical connection, which is an ideal test of whether DDC can “explain away” spurious correlations. We simulated both linear and sigmoid based nonlinear dynamics (Methods). We intentionally introduced a model mismatch - “sigmoid” based generative dynamics and “ReLU” nonlinearity for estimation - because it’s impractical to assume a perfect match. Both ΔL and ΔReLU correctly inferred the existence and direction of the ground truth connections (Fig. 2) while the covariance matrix (Cov) failed to explain away false positive connections and partial covariance (P) was not able to determine the directionality of connections (Fig. S2B).

**Fig. 2.**
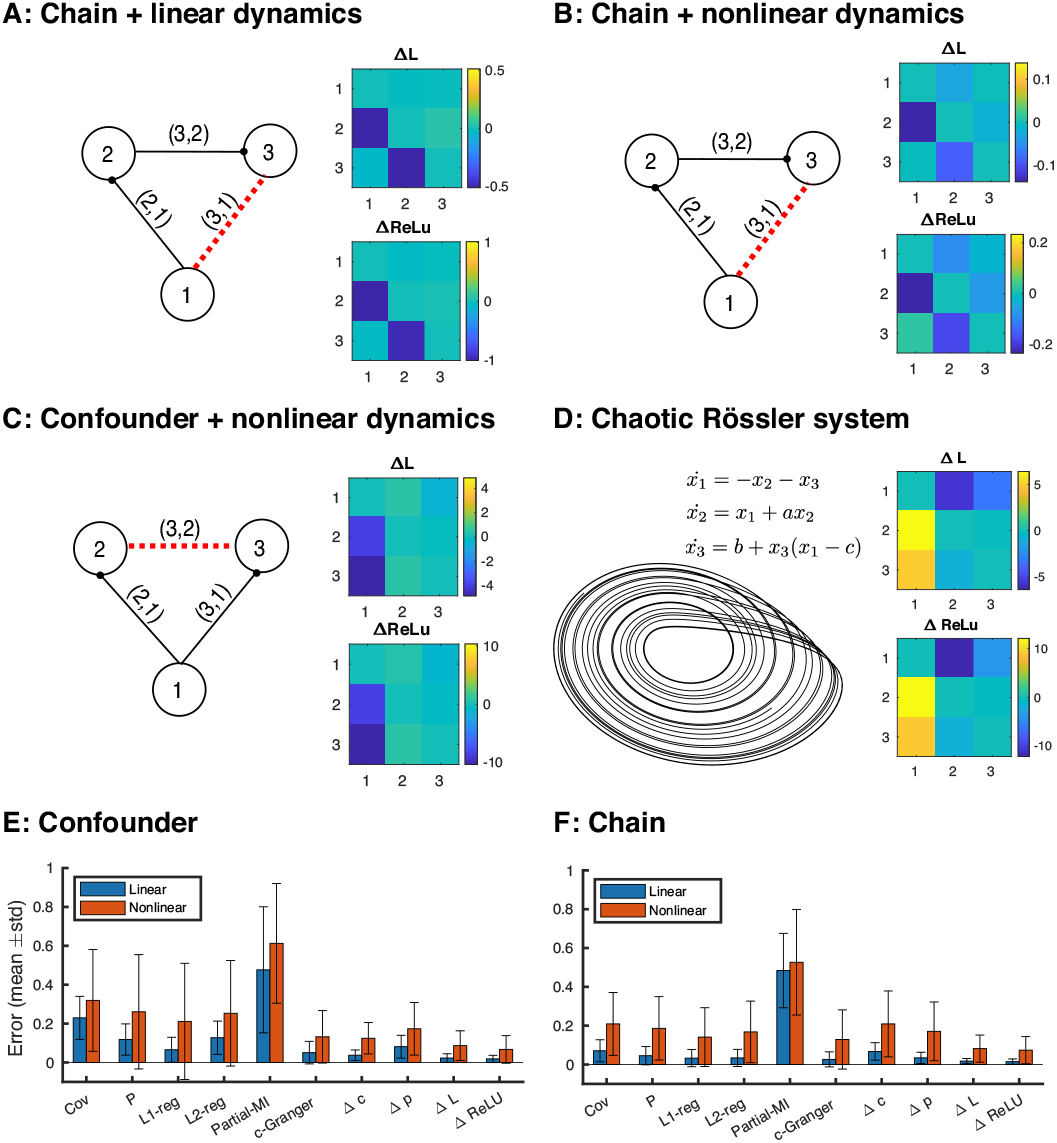
DDC recovers ground truth connectivity across multiple 3-node networks. (A)(B)(C) Left: ground truth network structure. Black solid lines are directed physical connections and red dashed lines are false positive connections commonly inferred by covariance estimation; The edges label (*i, j*) stands for the matrix entry at the *i*-th row and *j*-th column. Right: estimated ΔL and ΔReLU. (D) Left: phase diagram of *x*_1_ and *x*_2_ of the Rosseler system governed by system equations shown above; Right: estimated ΔL and ΔReLU. (E)(F) Estimation performance over 50 trials simulated confounder chain motif using both linear and nonlinear dynamics. L1/L2-reg: L1/L2-regularized partial covariance matrix; Partial-MI: partial mutual information; c-Granger: conditional Granger causality; std: standard deviation.

We further benchmarked DDC performance with more sophisticated methods, including regularized partial covariance, partial mutual information (MI) and conditional Granger causality (c-Granger) (Fig. 2E and F). L1 regularized partial covariance (L1-reg) and c-Granger exhibited comparable performance in linear simulations but their performance deteriorated for models with nonlinear dynamics. The heavy computation burden for optimization or model fitting makes these two methods difficult to scale up (See computation time in Fig. S2E for a 50-node task and in Fig. 3D for a 200-node task). Because regularized partial covariance and c-Granger assume stationarity, they performed poorly on the state-switching case shown in Fig. 1E and F.

**Fig. 3.**
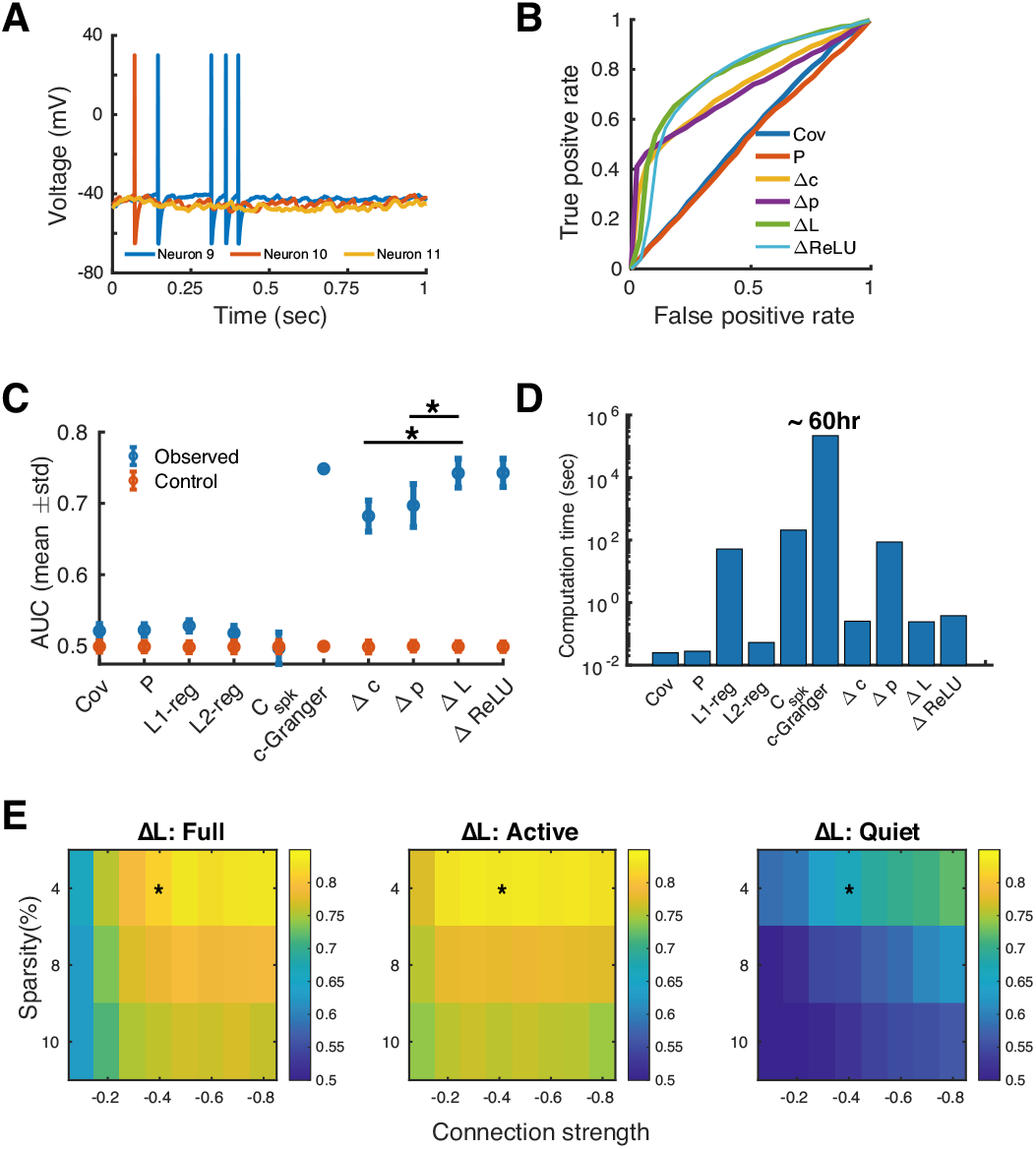
Estimation of performance for spiking networks: (A) Selected membrane potential traces simulated using LIF neurons with a sparse connectivity matrix. (B) Receiver Operating Curves (ROCs) quantifying classification performances for true connections for spiking networks of LIF neuron models. (network sparsity: 0.04, connection strength: -0.4, average firing rate: 3.5Hz). The curves for ΔL and ΔReLU were similar due to threshold selection process (Methods). (C) Area Under the ROC Curve (AUC) across 50 realizations of the random graph in (A) (*: p<0.001, two-sided Wilcoxon rank-sum test). Control values were calculated by randomly permutating the edges in the ground truth network. Only one AUC value was possible for the c-Granger estimate because of the excessive computation needed. C_spk_: spike train cross correlogram based connectivity. (D) MATLAB computation time on a computation cluster with 32 CPUs for the dataset of 200 nodes by 200,000 time points. (E) AUC values for ΔL applied to networks with a range of sparsities and connection strengths. Estimates used either (Full) all time points; (Active) time points around the spike times; or (Quiet) time points outside the spike time intervals. Asterisks indicate the condition used for ΔL in panel (C).

DDC was also applied to a larger network consisting of 50 nodes and structured by a combination of confounder and chain motifs (Fig. S2A). As in the small network case, ΔL and ΔReLU accurately estimated the existence and direction of connections (Fig. S2B). L1-reg and c-Granger achieved good performance for models with linear dynamics but performed poorly when the system was nonlinear (Fig. S2D). In this 50-node estimation task, c-Granger computation time was more than three order of magnitude greater than the others (Fig. S2E). For most methods, estimation performance improved with larger data sets (Fig. S2C).

Finally, we asked whether DDC can track the information flow in a nonlinear Rössler system, which is deterministic and chaotic. The three equations for this system in Fig. 2D have a nonlinear bidirectional confounder motif. ΔL and ΔReLU correctly identified direct connections, and ignored the strong correlations between *x*_2_ and *x*_3_. This suggests that DDC estimation is robust to model mismatch and can faithfully reflect the direct interactions in the system equation despite the unpredictability of the chaotic system.

### D. DDC identified ground truth connections with high sensitivity across multi-scale neural networks

Next, we tested DDC on a network model with 200 Leaky Integrate-and-Fire (LIF) neurons (21). These neurons integrate exponentially filtered synaptic inputs until the membrane potential reaches a threshold, which triggers a spike and a reset to resting membrane potential. The connectivity matrix was a globally connected Erdős–Rényi random graph with uniform connection strengths, emphasizing the existence and direction of network edges (Methods). Hence, we used the classification performance of true connections with increasing binarization thresholds (Methods) as a surrogate measure for the whole matrix estimation performance.

Graphs with a range of sparsity and connection strengths were simulated and DDC was applied to the subthreshold membrane potentials (Methods, representative traces shown in Fig. 3A). Performance was quantified by the area under the curve (AUC) of specificity versus sensitivity (Fig. 3B and C). Directed estimation methods (c-Granger, dCov and DDC) have higher AUC values because the ground truth matrices, which were not symmetric. ΔL and ΔReLU were significantly (p<0.001, rank-sum test) better than all other methods except c-Granger, which reached comparable performance level. However, one trial of c-Granger estimation took approximately 60 hours on a computing cluster with 32 cores, which makes it impractical to analyze a large number of neural recordings. Both the linear and nonlinear estimators presented similar curves (Fig. 3B) because the threshold selection process (Methods) of ΔReLU favored a low threshold, approximating a linear one. Possible explanations and improvements of the nonlinear estimator will be discussed later. Nonetheless, the linear DDC estimator remained its high performance in this highly nonlinear simulation and was robust to a broad range of network configurations with different sparsity levels and connection strengths (Fig. 3E). In general, the additional operation of taking partial covariance compared to pairwise covariance didn’t improve the performance (compare Cov, Δc with P, Δp in Fig. 3D) due to the sparsely connected network structure - the influence of indirect connection is trivial. Interestingly, Δp had very high sensitivity (true positive rate) even when very few connections were thresholded as positive. This might due to its sparse estimation (14) (16)

Due to the spike and reset process, the numerical derivative of membrane potential jumps transiently during spike events, thus significantly contributing to DDC estimators. We suspected the DDC estimators performed well because of these spike events. To test this possibility, we split the time trace and its derivative trace into active time points where at least one neuron spikes and quiet time points where no neuron emits a single spike. We extended the simulation to 40 seconds to ensure a sufficient data volume and quantified DDC performance for both parts (Fig. 3E, middle and right panel). Estimation of ΔL using active time points was as accurate as that using full time points. However, the spike train cross-correlogram was itself not enough to recover the connectivity as indicated by the low AUC value of C_spk_ in Fig. 3C and Fig. S3C). This indicated the importance of voltage fluctuations within the active period window, which is irreplaceable by binary spike train information.

We also tested the methods on simulated recordings of macroscopic neural activities based on the reduced Wong-Wang model of the resting state (22). The connectivity matrix used short-range local interactions and DSI (diffusion spectrum imaging) measurements of long-range structural connectivity (23) (Fig. 4A). The overall performance was quantified by C-sensitivity (Methods), which is a measure of the separation between the estimated value of true positive connections and the estimated value of true negative connections. C-sensitivity = 1 when the estimated matrix completely separated true positives from the others. ΔL had the highest performance followed by c-Granger and ΔReLU (Fig. 4B), probably because the reduced Wong-Wang model exhibited linear fluctuations around the stable point (22). Again, c-Granger exhibited comparable performance but the computation time was almost four order of magnitude longer, making it impractical. The raw ΔL and ΔReLU matrices uncovered the strongest connections (red arrows) in the ground truth matrix (Fig. S4B). (Only the strongest long-range connections were included, because all methods failed to reach significance for graded anatomical connectivity (Fig. S5).)

**Fig. 4.**
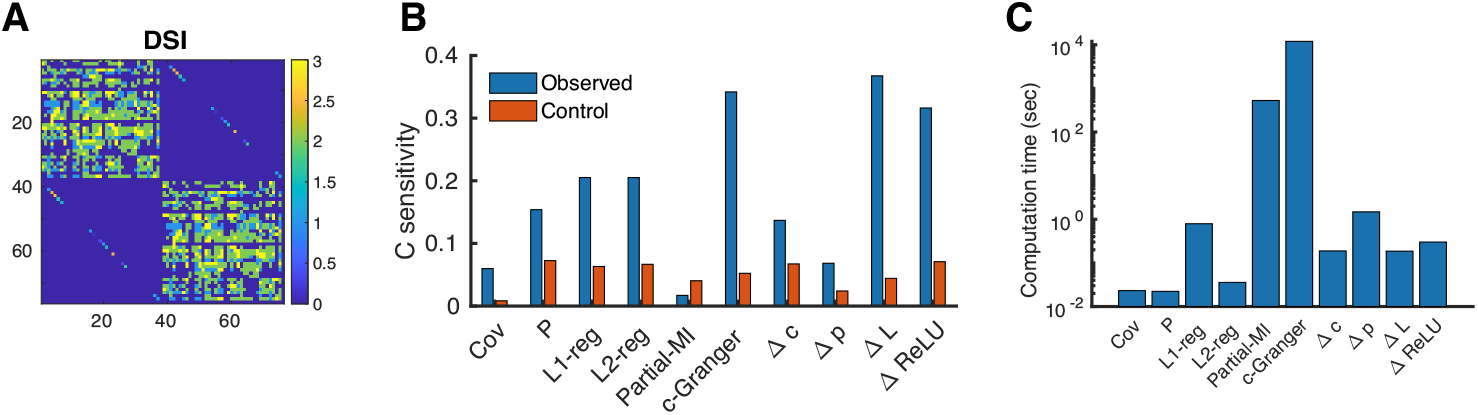
Estimation performance for a macroscopic brain surface model. (A) Long-range structural connectivity measured by Diffusion Spectrum Imaging (DSI). The block structure represents the left and the right hemispheres. (B) C-sensitivity of the estimators, a measure of the separation between estimated values of true positive connections and true negative connections. Control values were calculated by randomly permuting the edges in the ground truth network. (C) Computation time for the dataset of 76 nodes by 100,000 time points.

### E. DDC estimation is reliable when applied to rs-fMRI recordings

To critically test DDC on neural data, we applied DDC to resting-state fMRI (rs-fMRI) recordings obtained from the Human Connectome Project (HCP). The imaging voxels were parcellated through group ICA (Methods), where each independent component (IC) parcellation, shared across subjects, is composed of voxels with similar dynamics. ICs are mainly composed of spatially proximate voxels, forming anatomically recognizable brain regions (Fig. S8). In addition, we focused on the first 46 ICs encompassing over 40% cortical voxels (Fig. S7A) to match cortical measurements using diffusion MRI (dMRI). Dual regression (Methods) assigned unique ICA-parcellated BOLD signals to each subject, which were treated as nodes for DDC analysis.

We first established the reliability of ΔL estimation; that is, the number of data points required for estimates to become stable and whether the method provided consistent estimates across sessions within one subject. Within one subject, we calculated the correlation of ΔL estimation using all data points with that estimated using different amounts of data. The correlation value is 0.6 using data size equivalent to one recording session and gradually increases as more data are included (Fig. 5A). Compared to sample covariance estimation (Fig. S6), the data volume requirement is relatively higher probably due to the calculation of derivative. For most subjects, 2,400 data points, or equivalently, two sessions, were needed to reach a correlation of 0.8. We further plotted the correlation of ΔL estimation using a concatenation of two sessions within and between 10 randomly selected subjects (Fig. 5B). The block structure on the diagonal indicated a higher intra-individual similarity of ΔL across sessions. The population statistics from 1,003 subjects (Fig. 5C) confirmed a significantly higher within-subject similarity and a relatively lower (compared to Fig.S6C) between-subject correlation. Collectively, these results suggest that ΔL provides consistent estimates of brain connectivity within one subject, and at the same time preserves individual differences.

**Fig. 5.**
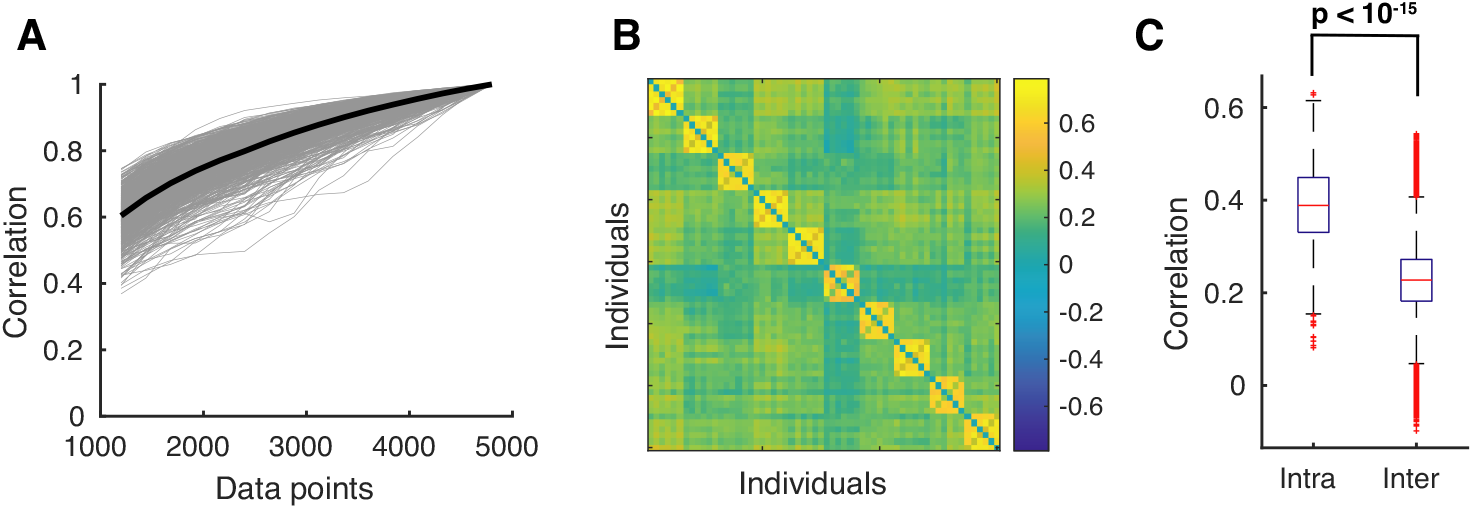
Intra- and inter-individual variability of ΔL estimation across scan session. A) Correlation of the full-length ΔL estimation with that estimated using different amount of data. One scan session includes 1200 data points. Each gray curve represents one subject and the black curve is the average. B) Correlation of ΔL estimated using a concatenation of two scan sessions within and between 10 randomly selected subjects. The block diagonal structure indicates a higher level of intra-individual similarity. C) Boxplot of intra- and inter-individual correlation pooled from all 1,003 subjects (two-sided Wilcoxon rank sum test, p< 10^−15^). The inter-individual correlation was only calculated between non-overlapping concatenation of sessions (i.e. 3 values per subject).

### F. DDC consistently identified structurally connected brain regions

The average and standard deviation of the estimated DDC matrices across subjects are shown in Fig. 6A and B where ΔL and ΔReLU were sparser than the covariance matrix. Two nodes (indicated by red arrow) in ΔL appeared to have a broader range of interactions. They were anatomically registered as “occipital pole” and “medial occipitotemporal gyrus”, reflecting the large proportion of visual ICs in the network (Fig. S8 and Table S1). To increase noise tolerance, we binarized the estimated matrices based on significance levels as determined by an auto-regressive bootstrapping procedure that preserved the signature power spectrum properties of BOLD signals (Methods). The significant ΔL connections (yellow entries, *p* < 0.01) across subjects were shown in Fig. 6C. These “backbone connections” shared across a majority of the subjects could be flexibly tuned for network sparsity level. We adopted a strict criterion because we were interested in the most conserved connections shared by over 90% subjects (red dashed vertical line in Fig. 6C). Their IC parcellations were registered on an MRI template (Fig. 6D). In this case, “backbone connections” were identified between ICs from the same anatomical region as well as inter-regional interactions (marked in red).

**Fig. 6.**
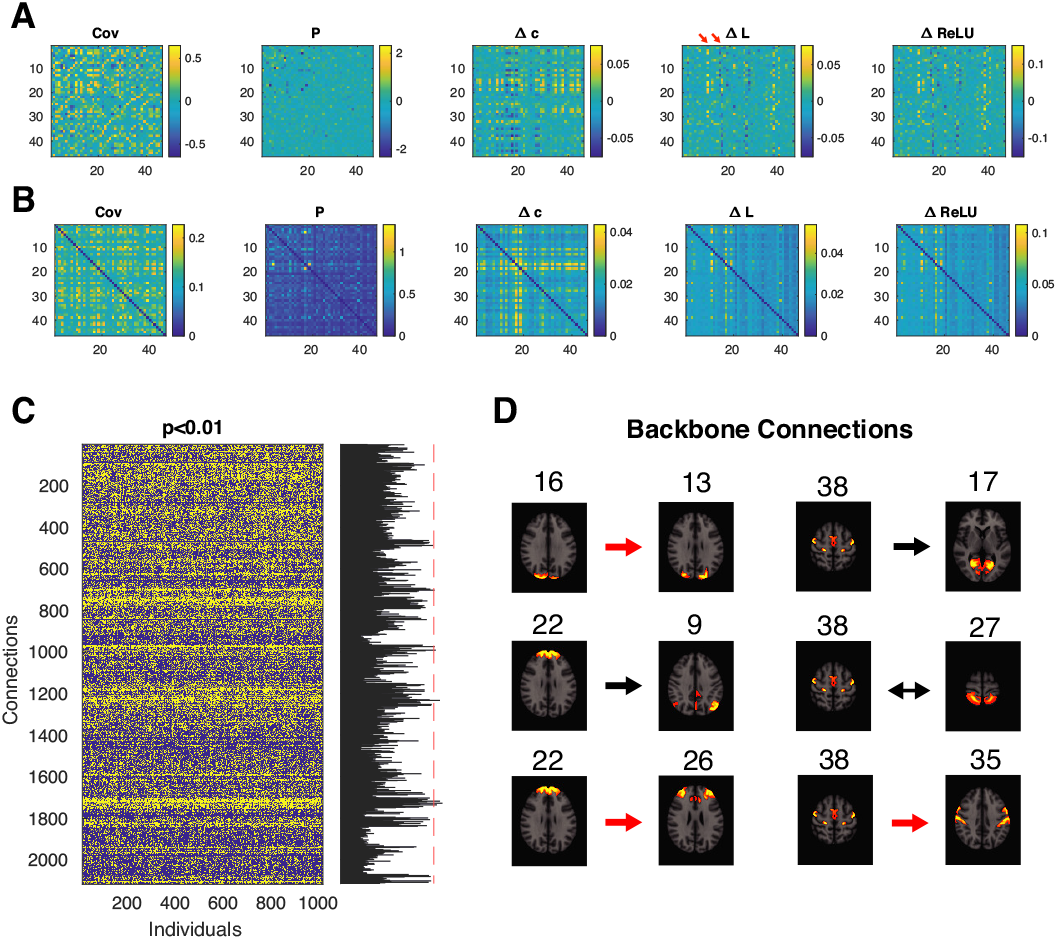
DDC consistently recovered known connections across HCP subjects. (A)(B) Average/Standard deviation of estimated FC matrices averaged across HCP subjects. (C) Significant (p<0.01, yellow entries) ΔL connections for 1003 individuals. Each column is a binarized matrix reshaped to a column vector. The summation of each row, shown on the right, represents the number of individuals that highlighted a specific connection. “Backbone connection” are those shared by most individuals (example threshold indicated by the red dashed line). (D) ΔL “backbone connection” shared by over 90% subjects and their IC parcellations registered on an MRI template. Arrows indicated the estimated connection direction and the red arrows indicate IC pairs that are anatomically close. Numbers indicate the IC indices corresponding to Table S1 and Fig. S8

To quantify the extent to which estimated FCs matched the structural connectivity, we further processed dMRI measurements from the HCP dataset (24) to obtain individual-level IC-based dMRI matrices (Fig. S7, Methods). At the IC level, dMRI strengths were bimodal (Fig. 7A), indicating a clear separation between the strong and weak connections. ΔL identified connections with higher dMRI strength compared those chosen by the covariance matrix (Fig. 7B). Fig. 7C shows the increasing average dMRI strength for decreasing binarization threshold, linking the significance of rs-fMRI to dMRI connectivity for all methods and confirming their biological relevance. DDC uncovered connections with significantly higher dMRI strength values than covariance-based methods (Fig. 7C) and also identified a larger proportion of strong connections (Fig. 7D).

**Fig. 7.**
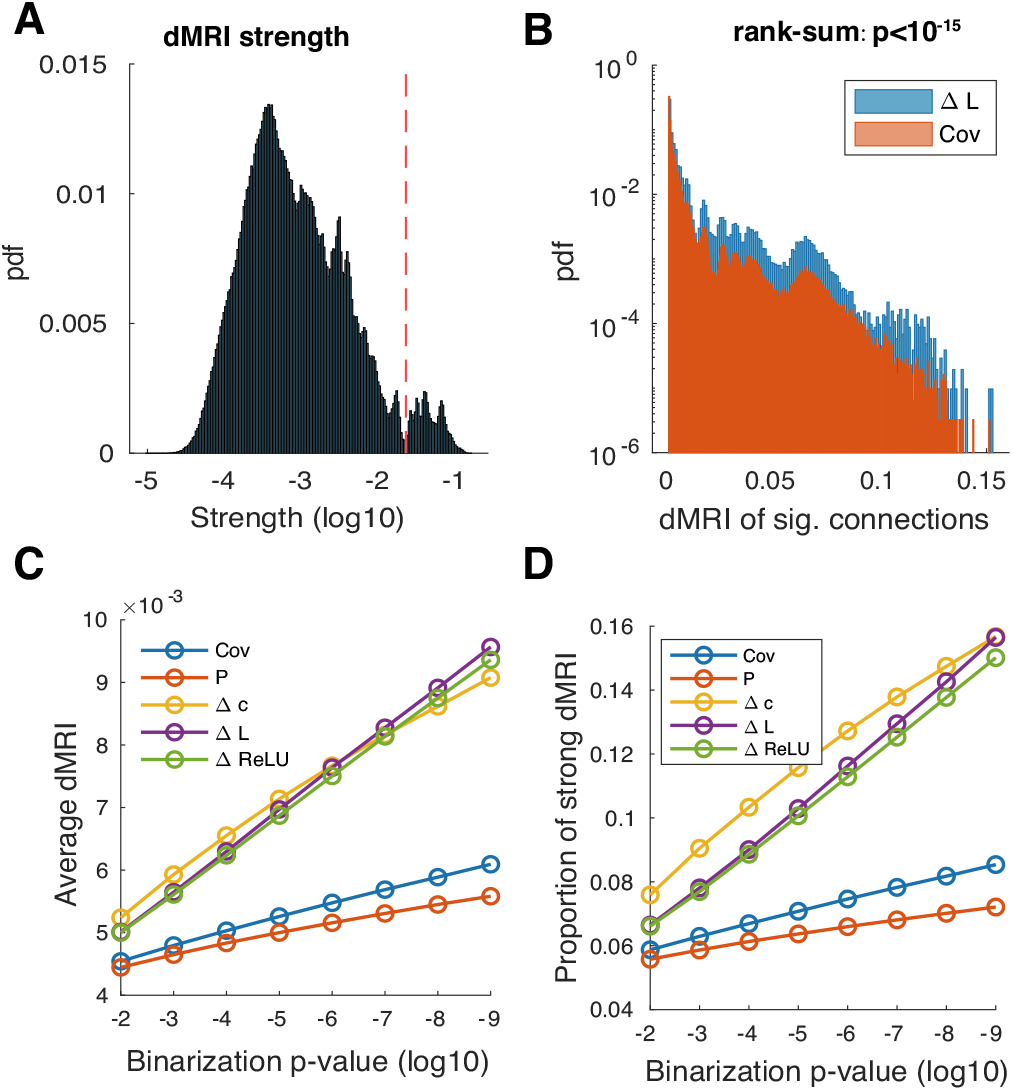
DDC picked up connections with strong dMRI values. (A) Distribution of IC-level dMRI strengths. Connections to the right of the cutoff value (red dashed line) were classified as strong connections. (B) The dMRI strength distribution of significant ΔL and Cov connections (Binarization p-value< 10^−9^; two-sided Wilcoxon rank sum test, p< 10^−15^). Note the log scale on the y axis presented due to the large abundance of weak connections. (C) Average dMRI strength value of significant connections picked by different methods with stricter binarization thresholds. (D) Proportion of strong connections. Since dMRI strength distribution is bimodal, the proportion of strong connections was also used as a supplementary statistics to compare the distributions.

## 3. Discussion

Dynamical Differential Covariance is a promising new family of estimators for analyzing the functional connectivity underlying large-scale brain recordings. Because DDC is derived directly from dynamical system equations that govern neural interactions, no optimization or model fitting is required. DDC is a practical and intuitive method that can be computed rapidly and scales well with the number of recording sites. Unlike methods based on covariance, which are inherently symmetrical, DDC can detect directional interactions and obtain statistical estimates of causality.

DDC uncovered ground truth when applied to dynamical simulations of network models and significantly improved estimates of strong dMRI connectivity from rs-fMRI recordings compared with covariance methods. Further analysis is needed to probe the ability of DDC methods to uncover weaker connectivity. Structural anatomy is foundational in neuroscience and functional anatomy that is consistent with the underlying structural anatomy makes possible more accurate predictions about information flowing in specific pathways.

There are many ways that DDC can be extended, particularly in the choice of the underlying model of the dynamics that generates the neural activity. Here, we adopted a ReLU nonlinearity, which has been used to model nonlinear rate-based network models, but its performance on the spiking network model was no better than the linear model. A better model for a spiking network would take into account the spiking mechanism and dynamical time constants. See the Materials and Methods G.3 for further discussion on how to modify nonlinear DDC for spiking networks.

In conclusion, DDC has a number of favorable mathematical properties that should ensure robust estimation of FC from a wide range of noisy and nonstationary recordings. Access to the directionality of neural connections opens new avenues for interpreting the causal flow of information through networks. Identifying functional connectivity based on dynamical systems models makes direct contact with similar approaches in many other disciplines such as bioengineering, control theory and network science. DDC should have a broad impact on studies in these areas whenever there is need for estimating directional network connectivity from network activity.

## ACKNOWLEDGMENTS

We are grateful to Tiger Wutu Lin, Anup Das, Qasim Bukhari, Maksim Bazhenov, Syd Cash and Eric Halgren for previous collaborative research and for helpful discussions. The authors also thank Jorge Aldana for assistance with computing resources. This research was supported by the Office of Naval Research (N00014-16-1-2829, N00014-16-1-2415), NIH/NIBIB (R01EB026899), and NIH/NIMH (1RF1MH117155-01). Human rs-fMRI and dMRI datasets were provided by the Human Connectome Project, WU-Minn Consortium and the McDonnell Center for Systems Neuroscience at Washington University.

## Data and materials availability

Implementation of DDC, network simulation and HCP processing scripts are all available through https://github.com/yschen13/DDC.

## Supporting Information Appendix (SI)

See the SI pdf for Table S1 and Figure S1-S8

## Materials and Methods

### 1. Functional Connectivity Estimators

All estimators and abbreviations are summarized in Table 1.

**Table 1.**
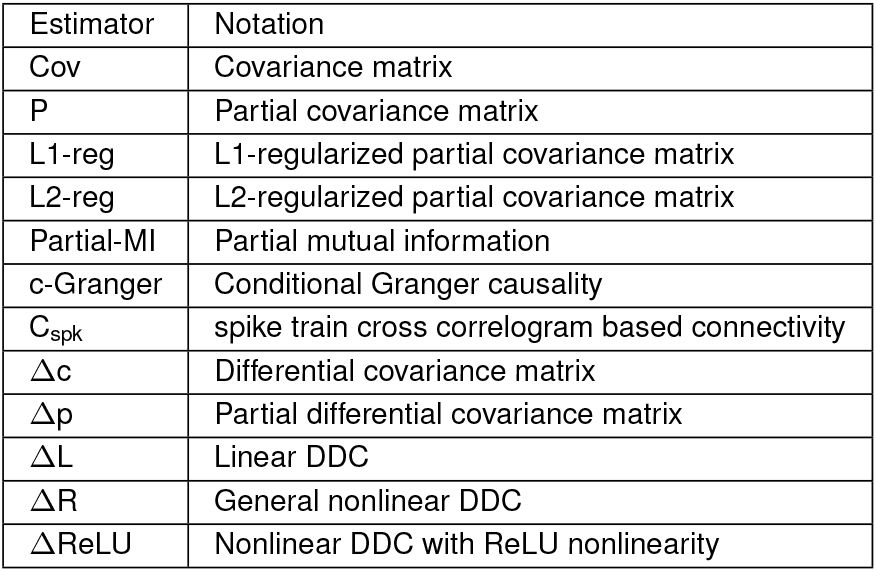
Summary of estimators.

| Estimator | Notation |
| --- | --- |
| Cov | Covariance matrix |
| P | Partial covariance matrix |
| L1-reg | L1-regularized partial covariance matrix |
| L2-reg | L2-regularized partial covariance matrix |
| Partial-MI | Partial mutual information |
| c-Granger | Conditional Granger causality |
| $C_{\text{spk}}$ | spike train cross correlogram based connectivity |
| $\Delta c$ | Differential covariance matrix |
| $\Delta p$ | Partial differential covariance matrix |
| $\Delta L$ | Linear DDC |
| $\Delta R$ | General nonlinear DDC |
| $\Delta \text{ReLU}$ | Nonlinear DDC with ReLU nonlinearity |

#### A. Covariance based estimators

The covariance (Cov) and partial covariance (P) matricies are:

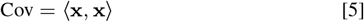

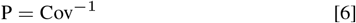

where **x** is a column vector of the system variable and the operation ⟨, ⟩ takes the outer product of two vectors and averages across time. In this paper, all time traces were z-scored, thus, the covariance matrix is equivalent to correlation. The covariance matrix only reveals pairwise correlations but the partial covariance matrix controls for confounding effects, one step closer to a causal estimation of global connectivity.

#### B. Regularized partial covariance

Under the assumption that the connectivity matrix is sparse, regularization methods could be implemented during the regression step of calculating partial covariance matrix. For example, we calculated L1- and L2-regularized partial covariance matrices through the FSL toolbox (25). To choose the regularization parameter (*λ*), we tested a range of them and chose the one with the best performance with corresponding quantification metric. We tested *λ* = 5, 10, 20, 50, 100, 200 for L1-regularization while *λ* = 0.1, 0.2, 0.5, 1, 5 for L2-regularization.

#### C. Mutual information

Mutual information (MI) quantified the statistical dependence of two random variables beyond second-order statistics. It’s a model-free estimation of dependencies therefore it should work equally well for both linear and nonlinear simulation. We used the implementation in the Functional Connectivity Toolbox (26) to estimation Partial MI since we want to estimate the global network structure.

#### D. Granger causality

Granger causality (8) defines a statistical interpretation of causality based on predictability: A is said to “Granger cause” B if the predictability of B declines when A is removed from the predictors.The test of predictability increase or decline is usually implemented through multivariate vector auto-regressive modeling. We implemented conditional Granger causality through the MVGC MATLAB toolbox (27). In this approach, the fundamental assumption is that the time series data is a realization of a stationary vector auto-regressive (VAR) process. The VAR model order was chosen based on Akaike information criterion (AIC) and the coefficients of the full/reduced regression model was computed through Ordinary Least Square (OLS) solution to the regression.

#### E. Spike train cross correlograms

Following Das et al (28) and Guisti et al (29), we calculated the cross correlograms based on their Pearson correlation and averaged within a time window (*τ* = 0.1 sec) to get a spike based connectivity measure *C*_spk_. Specifically, for spike train *x*_*i*_ and *x*_*j*_, 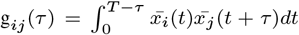 where the superscript bar denotes the centered version of the spike train. Then the connectivity value from *j* to *i* was evaluated as:

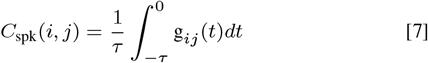

For numerical calculation, we used the sampled time interval (dt = 0.5 msec) to evaluate the integral. For neuron pairs whose firing rates are smaller than 0.1 Hz, their connectivity value was assigned as zero. Around 2,000 connection pairs were skipped in Fig.S3C.

#### F. Differential Covariance (dCov) estimators

Differential covariance (Δc) was calculated as Eq.8 where 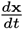 was numerically computed using a symmetric difference quotient (30). The evaluation of partial differential covariance (Δp) was derived in parallel to partial covariance. The calculation was performed element-wise as in Eq.9 where Cov refers to the covariance matrix, *K* denotes the set of all nodes except *i* and *j*.

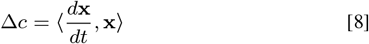

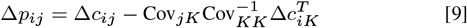

#### G. Dynamical Differential Covariance (DDC)

The definitions of ΔL, ΔR and ΔReLU can be found in the main text. The parameter *θ* for ΔReLU, was varied from the 5th percentile to the 95th percentile of the z-scored data. The optimal value was chosen based on either the estimation errors (three-neuron simulations) or the AUC values (LIF neuron simulations). In the brain surface model, *θ* was set to zero.

##### G.1. DDC derivation for stochastic network models

To model the randomness in the recorded neural activities, we used stochastic differential equations (SDE) and evaluated DDC in this stochastic framework:

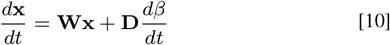

where *β* is a multi-dimensional Brownian motion with variance **Q** and noise structure **D** influencing the state variable **x**. See Eq. 1 in the main text for the definitions of the other terms. The time averages 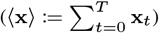 are different from ensemble averages (𝔼 (**x**)) under nonstationary conditions, as analyzed in the next section.

Operating on both sides of this equation with ⟨, **x**⟩:

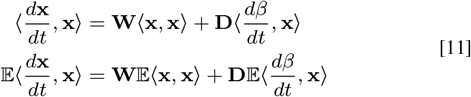

To evaluate 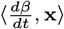, we first write down the explicit solution of the linear SDE starting at *t* = 0, then time average both sides.

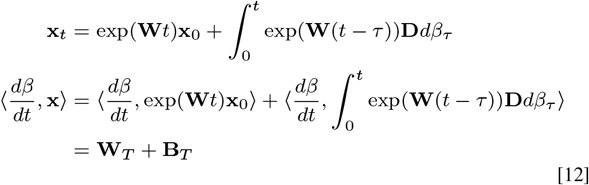

The first term **W**_*T*_ is the summation of time-dependent linear Brownian increments, thus the mean is zero and the variance is a time-dependent scaling of the Brownian variance:

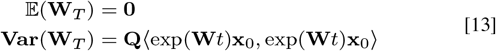

The second term **B**_*T*_ was evaluated using the Ito integral. Because Brownian motion is nowhere differentiable on its path, we numerically approximated the time derivative as we used in the simulations. If we assume 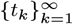 as a partition of [0, *t*] whose partition size is infinitesimal as *n* → ∞, we can compute Ito integral in the limit. For simplicity, define **Φ**_*τ*_ = exp(**W**(*t* − *τ*))**D**. Then the two terms are composed of nonoverlapping Brownian increments:

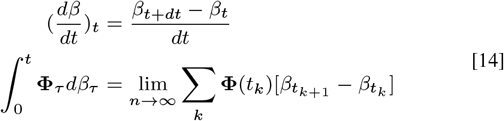

from which it follows that 𝔼 (**B**_*t*_) = 0 because Brownian motion has independent and stationary increments.

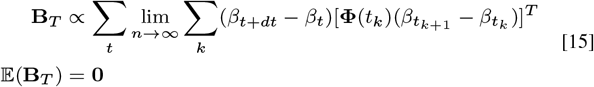

Taken together, the first order statistics of our linear DDC estimator become:

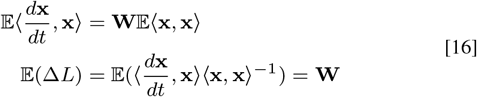

This derivation confirms that DDC is unbiased in the presence of noise as a linear combination of Brownian motion. Simulations of the linear three neuron model revealed that 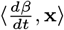 is at least ten times smaller in magnitude than ⟨**x, x**⟩ even for very high noise variance **Q**. Remarkably, DDC can still recover the ground truth connectivity for correlated noise structure (**D** in Eq.10).

##### G.2. Nonstationary conditions

A continuous time stochastic solution of the SDE, 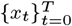, is stationary when its finite-dimensional joint distribution is time-invariant, which implies that its mean and covariance remain constant across time. Only under the stationary assumption, the covariance matrix can then be estimated by the time-averaged sample covariance.

The above SDE framework allows the mean and covariance of state variables to vary with time according to the Ito formula:

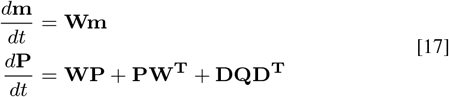

where **m** = 𝔼 (**x**), **P** = **Var**(**x**). The process is stationary if the right hand sides are zero. The steady state solution of **m** is either zero or within the null space of **W**. Meanwhile, the steady-state solution of **W** is given by the Lyapunov matrix equation and can be solved by vectorization (*v*) and Kronecker product (⊗):

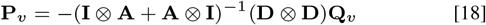

Under nonstationary conditions, ⟨ **x, x** ⟩ is no longer a valid estimate of the covariance matrix. Because the system equation holds at every time step regardless of stationarity, our DDC estimators remain valid and unbiased. These properties make DDC a robust and efficient estimator of functional conectivity.

##### G.3. Modifications of the nonlinear estimator

Models of spiking neurons have several features that are not included in the models studied in this paper. First, decay time constants for neurons and synapses are an important part of their dynamics, such as the leaky dynamics of LIF models. Consider the following linear and nonlinear system equations that include membrane time constants for temporal filtering:

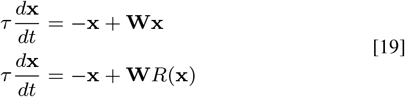

For the linear case, DDC estimation only differs up to a scaling factor and the diagonal terms:

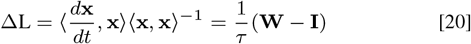

For the nonlinear system, the time decay introduces a new term:

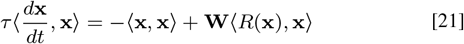

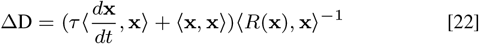

where ΔD is a different DDC estimator of **W** for the temporally-filtered nonlinear system. It is proportional to a weighted average of dCov and Cov. The time constant in this equation is a free parameter that can be estimated or optimized. This approach allows for a better match with the circuit mechanisms that generate the activity and how that activity is transformed by the recording techniques.

A second modification to DDC concerns the sparsity of the spiking. For the simulated LIF network, the sparse spike trains were filtered with decaying synaptic currents. As a consequence, ⟨*R*(*x*), *x* ⟩ was rank-deficient and thus non-invertible. One potential remedy is to find the least square solution of a sequence of linear equations to optimize **W**. This will be explored in a future study.

### 2. Simulations of neural systems

#### A. Neural motif dynamics

We tested the performance of these methods in networks structured to have typical false positive motifs - chain (Fig. 3A,B) and confounder (Fig. 3C) - with different dynamics and in another Rosseler chaotic system. To stabilize the simulation, all nodes had decaying dynamics and they were linked by inhibitory connections. Specifically, the diagonal entries in the ground truth matrix was set to -1 and connection strength was set to -0.5. We tested connection strength from -0.1 to -1 and it didn’t affect the estimation results qualitatively.

For linear dynamics, system variable **x** was simulated through Euler integration according to Eq.23 where **u** the is Gaussian distributed random drive and *ϵ* is the Gaussian distributed observational noise, both independent from **x**. The integration step is 0.01 seconds and the length of simulation is 1000 seconds unless otherwise specified.

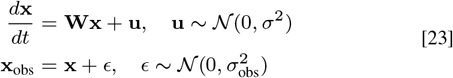

For nonlinear dynamics, simulation was governed by Eq.24 where we used a centered sigmoid function to simulate the nonlinearity. The sigmoid function was shifted to have mean of zero because otherwise the inhibitory signal would be too strong in the network and the signal would decay to zero in a short time interval. In the expression of *R*(*x*), slope *α* controls the level of nonlinearity in the network and was set to 1 by default. Note the mismatch between simulation nonlinearity and the estimation nonlinearity. The integration step is 0.1 ms and signals were downsampled to 100 Hz after estimation. Simulation length is 1000 seconds unless otherwise mentioned.

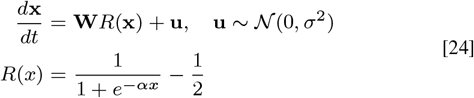

The equations for the Rössler system are:

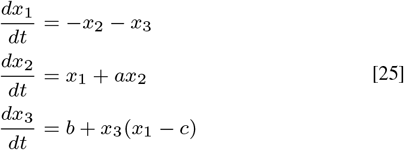

where **x** = [*x*_1_, *x*_2_, *x*_3_]^*T*^, *a* = *b* = 0.2 and *c* = 5.7. This set of parameters was originally used by Rössler to study the behavior of its chaotic dynamics. The signal was integrated at the step of 0.01 seconds for 1000 seconds. The first 100 seconds of transient dynamics was discarded.

To simulate a nonstationary system with time-invariant connectivity pattern, we borrowed the idea of hidden markov models and simulated a two-state dynamical system. The system was simulated for 1000 seconds using Euler integration governed by Eq.26. Up until 500 seconds, the noise stucture remained to be the identity matrix. After 500 seconds, we switched the noise structure to **D**_**2**_. The steady-state covariance structure of these two states could be analytically evaluated using Eq.18 and plotted in Fig. 2A.

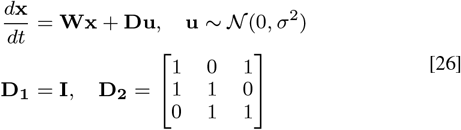

#### B. Sparse Leaky Integrate-and-Fire (LIF) network

The connectivity matrix was constructed as an Erdős–Rényi random graph: two nodes being connected has probability equal to network sparsity. All connected edges were assigned to have the same strength. Thus, the connectivity matrices were parameterized by only sparsity level and connection strength. Leaky-Integrate-Fire neurons could be described by Eq.27 with double-exponential filtered synapses (21). Once membrane voltage *V* reaches a threshold *V*_*thres*_, the neuron will emit a spike and reset the membrane potential to *V*_*reset*_. The spike train was described by 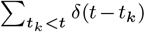 and then filtered to generate synaptic current *r*_*i*_. We used sub-threshold membrane potential as the system variable (**x**) of interest. We simulated networks with 200 neurons. The integration process was performed at the step of 0.05 ms, down-sampled to 2000 Hz and simulated for 20 seconds unless otherwise mentioned.

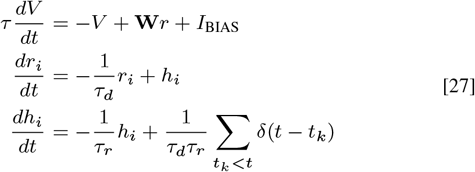

#### C. Anatomically supported brain surface model

The anatomic connection used here is the group average structural connectivity obtained through diffusion spectrum imaging (DSI) (23). It involves 78 cortical regions from both hemispheres. The connectivity matrix contained both local proximity based connections and long-range DSI measured connections. It was simulated using a reduced Wong-Wang model (Eq.6-8 in (22)) using the Virtual Brain Simulator (31). All physiological parameters were followed from the reference (22). Simulated population firing rates were down-sampled to 1000Hz and the simulation length was 100 seconds.

### 3. Estimator performance quantification

#### A. Variance and bias

Following Das et al (28), we decomposed the estimation error into variance and bias (Fig.S1). In most cases, the estimation is different from the ground truth matrix by a scale. So we normalized both estimated and ground truth matrices between -1 and 1. In addition, dCov based estimators are directed estimators while covariance based ones are not. For fair comparison, we only considered the estimation of the lower triangle part where all ground truth connections are located.

After scaling and lower triangle restriction, estimation error, variance and bias were calculated as Eq.28 where **W, Ŵ** and 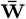 are ground truth matrix, estimated matrix and the average of estimated matrices across trials and ‖. ‖ is the vector L2 norm. It’s easy to verify that Error^2^ = Bias^2^ + Variance^2^ and thus the vector form of bias and variance are orthogonal to each other. We measure the relative contribution of bias by the angle (*θ*_*b*_, Eq.29) between the vectors associated with bias and variance. 50 repetitive trials were used across all simulations.

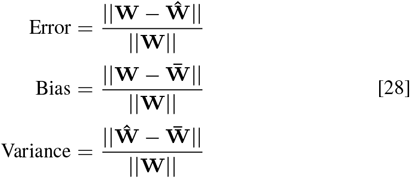

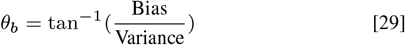

#### B. Sensitivity and specificity

To evaluate the estimation performance in LIF networks, connection recovery sensitivity and specificity were calculated since the networks have sparse connection and the connection strength are uniform. To be more specific, the estimated matrices were binarized based on their absolute values to determine the existence of connections, which were then compared with the ground truth connections. We used the absolute value because we only cared about the presence of a connection. Sensitivity and specificity were calculated as the true positive rate and one minus false positive rate. Varying the binarization threshold gave rise to the receiver operator curve (ROC). The area under ROC, calculated by trapezoidal integration, indicates the methods general performance in classifying connections.

For performance evaluation in the brain surface model, c-sensitivity (1) (Eq.17 in (15)) was adopted. It’s defined as the fraction of the estimated true positives values that are higher than the 95th percentile of the false positive values. Like ROC, c-sensitivity quantitatively estimated how sensitive methods are to estimating the presence of a connections. Thus, the absolute value of the estimated matrices were used here.

### 4. HCP dataset

#### A. Extracting time traces from rs-fMRI recordings

We used the extensively processed “HCP1200 Parcellation + Timeseries + Netmats (1003 Subjects)” dataset available through the website (https://www.humanconnectome.org). Detailed pre-processing and study design could be easily accessed through the website. In this release, 1003 healthy adult human subjects (ages 22-37 years, 534 females) were scanned on a 3-T Siemens connectome-Skyra scanner (customized to achieve 100 mTm^−1^ gradient strength). Each subject underwent 4 × 15 minutes recording sessions with temporal resolution of 0.73 second and spatial resolution of 2 mm isotropic.

For imaging data processing, each 15-minute run of each subject’s rs-fMRI data was pre-processed according to Smith et al (32); it was minimally-preprocessed (33), and had artefacts removed using ICA+FIX (34) (35). Inter-subject registration of cerebral cortex was carried out using areal-feature-based alignment and the Multimodal Surface Matching algorithm (‘MSMAll’) (36) (37). Each dataset was temporally demeaned and had variance normalization and then fed into the MIGP algorithm, whose output is the top 4,500 weighted spatial eigenvectors from a group-averaged PCA (a very close approximation to concatenating all subjects’ timeseries and then applying PCA) (38). The MIGP output was fed into group-ICA using FSL’s MELODIC tool, applying at several different dimensionalities (D = 25, 50, 100, 200, 300). In our analysis, we used the 100-dimension decomposition.

For a given parcellation (group-ICA map), the ICA spatial maps were used to derive one representative time series per IC per subject. This process was fulfilled by the standard “dual-regression stage-1” approach, in which the full set of ICA maps was used as spatial regressors against the full data (39). This results in an *N* × *T* (number of components × number of time points) matrix for each subject. Thus, we consider each IC as a network node.

#### B. Intra- and inter-individual variability

The intra-individual variability was quantified as the correlation between DDC estimations across scans. Since DDC requires data points from at least two scans to achieve a relatively high consistency, we concatenated data from two scans for DDC estimation and compared it with the estimation results from the other two remaining scans (Fig. 5C). For inter-individual variability, estimation results of all six different combinations of concatenation were compared.

#### C. Significance test of the estimated connections

To assess the statistical significance of the estimated connection, we used an autoregressive (AR) bootstrap procedure (40, 41) to preserve the power spectrum density (PSD) of BOLD signals. For a specific estimated connection, denoted as element (*i, j*), our null hypothesis was that signal *x*_*i*_ and *x*_*j*_ are independent regardless of other nodes’ influence. To generate null time series, we fit separate AR processes of model order *q* to node-specific time traces. The model order *q* was determined according to the Bayesian information criterion (BIC). A higher order model was rejected if it could not decrease BIC by more than 2. Using the estimated AR coefficients of empirical time series, we generated 1000 surrogate null time series and then computed the associated functional connectivity corresponding to the null hypothesis. For each connection, we assumed a Gaussian distribution of the null connectivity values generated from null time traces. P value was calculated as the probability of the empirical value appeared under the null Gaussian distribution. In this paper, we adopted a sequence of significance level to binarize the matrix so that we could investigate the network behavior asymptotically.

#### D. Individual level dMRI strength

In order to compare the functional connectivity metrics to the underlying corticocortical white matter connectivity, we reorganized our previously published diffusion-MRI based structural connectome (24) in which connectivity was assessed among the 360 cortical areas of the HCP-MMP1.0 atlas (37). Of the 100 IC nodes, 46 are composed of at least 40% cortical voxels (Fig.S7A) and as the dMRI connectome was restricted to corticocortical relationships, we limited the scope of our analyses to these nodes. Because the IC nodes have a greater spatial extent than the atlas areas, each is composed of several areas, in whole or in part (mean = 28.3 areas). For each IC node pair, dMRI connectivity was assessed by obtaining the average of the nodes’ constituent interareal connectivity weighted by fraction of the node pair’s voxels assigned to each areal pair. In cases where an atlas area was partially present in both IC nodes of a pair, that area was excluded from the mean as short-range intra-areal anatomical connectivity was not available.

