## Supplementary material for "Dynamical differential covariance recovers directional network structure in multiscale neural systems": Fig S1-S8; Table S1

### This PDF file includes:

Figs. S1 to S8
Table S1
SI References

**Table S1. Anatomic annotations of the 46 ICs. First column is the index number of ICs. Second column is the manually registered anatomical region. DMN: default mode network; NA: no reasonable region identified**

| Component number | Anatomical Region | Sub-network |
|---|---|---|
| 1 | Occipital Pole | Visual Network |
| 2 | Inferior Parietal Lobe | DMN |
| 3 | Lateral Occipital Cortex | Visual Network |
| 4 | Cuneal Cortex/Occipital Pole | Visual Network |
| 5 | Supramarginal Gyrus | DMN |
| 6 | Lateral Occipital Cortex | Visual Network |
| 7 | Supramarginal Gyrus | DMN |
| 8 | Lateral Occipital Cortex | Visual Network |
| 9 | Inferior Parietal Lobe | DMN |
| 10 | Medial Prefrontal Cortex/Anterior Cingulate Cortex/Lateral Temporal Cortex | DMN |
| 11 | Lingual Gyrus/medial occipitotemporal gyrus | Visual Network |
| 12 | Angular Gyrus | DMN |
| 13 | Occipital Pole | Visual Network |
| 14 | Lateral Occipital Cortex - Left | Visual Network |
| 15 | Precuneous Cortex | Other Network |
| 16 | Occipital Pole | Visual Network |
| 17 | Lingual Gyrus/medial occipitotemporal gyrus | Visual Network |
| 18 | Lateral Occipital Cortex | Visual Network |
| 19 | Occipital Pole | Visual Network |
| 20 | Inferior Parietal Lobe | DMN |
| 21 | Precentral Gyrus | sensorimotor network |
| 22 | Orbital Frontal Cortex | DMN |
| 23 | Postcentral Gyrus | sensorimotor network |
| 24 | Lateral Occipital Cortex - Right | Visual Network |
| 25 | Occipital Pole - Left | Visual Network |
| 26 | Frontal Pole | Attention Network |
| 27 | Superior Parietal Lobule | Attention Network |
| 28 | Hippocampus/Parahippocampal Cortex | DMN |
| 29 | Lateral Temporal Cortex | DMN |
| 30 | Lateral Occipital Cortex | Visual Network |
| 31 | Orbital Frontal Cortex/Lateral Temporal Cortex | Other Network |
| 32 | Occipital Pole | Visual Network |
| 33 | Inferior Parietal Lobe – Left | DMN |
| 34 | Lateral Occipital Cortex | Visual Network |
| 35 | Postcentral Gyrus | sensorimotor network |
| 36 | Medial Prefrontal Cortex/Anterior Cingulate Cortex | DMN |
| 37 | Orbital Frontal Cortex/Lateral Temporal Cortex | DMN |
| 38 | Precentral Gyrus / Juxtapositional Lobule Cortex (formerly Supplementary Motor Cortex) | sensorimotor network |
| 39 | Occipital Pole | Visual Network |
| 40 | Middle Frontal Gyrus | Attention Network |
| 41 | Postcentral Gyrus - Left | sensorimotor network |
| 43 | Postcentral Gyrus - Right | sensorimotor network |
| 44 | Orbital Frontal Cortex | DMN |
| 45 | Superior Temporal Gyrus | sensorimotor network |
| 47 | NA | NA |
| 48 | Frontal Pole | Attention Network |

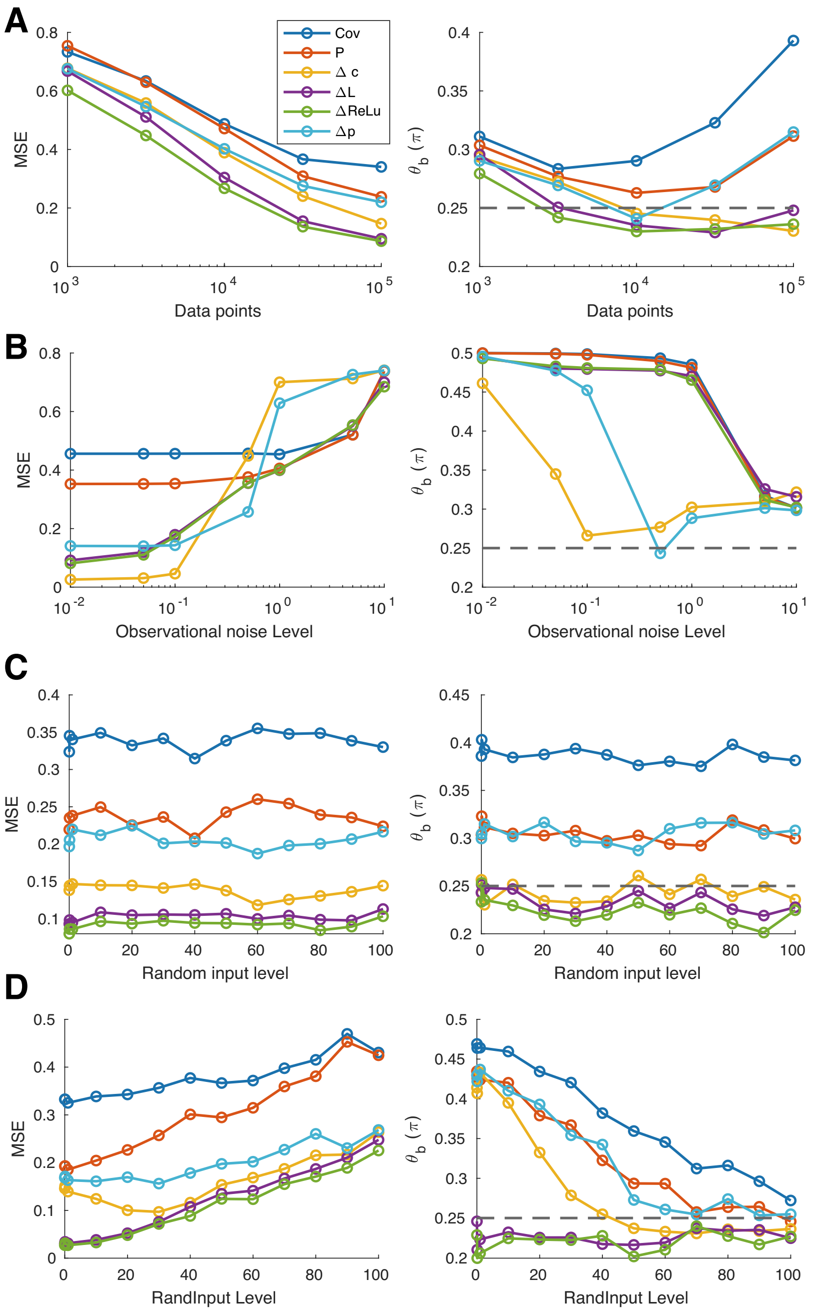


**Fig. S1. Estimator variance and bias under different parameter configurations** (A) Left: Influence of data volume calculated across 50 trials simulated using linear dynamics and the confounder motif; Right: Contribution of estimation bias ($\theta_b$). $\theta_b = 0.25\pi$ refers to equal bias and variance. In general estimation errors decreased with data volume and covariance based estimators are more biased. (B) Influence of observational noise (horizontal axis is $\sigma_{\text{obs}}$ in Eq.23) across 50 trials simulated using linear dynamics and the confounder motif. $\Delta$c and $\Delta$p performance deteriorated with larger amount of noise while $\Delta$L and $\Delta$ReLU remained robust under the influence of observational noise. (C) Influence of random input strength (horizontal axis is $\sigma$ in Eq.23) across 50 trials simulated using linear dynamics and the confounder motif. Input strength had limited effect on estimator performance since DDC is an unbiased estimator. (D) Influence of random input strength across 50 trials simulated using highly nonlinear dynamics ($\alpha$=50 in Eq.24) and the confounder motif. In general, the higher the input strength, the worse the estimation performance. Note the superior performance of $\Delta$L and $\Delta$ReLU over all other estimators when the random input strength was relatively low.

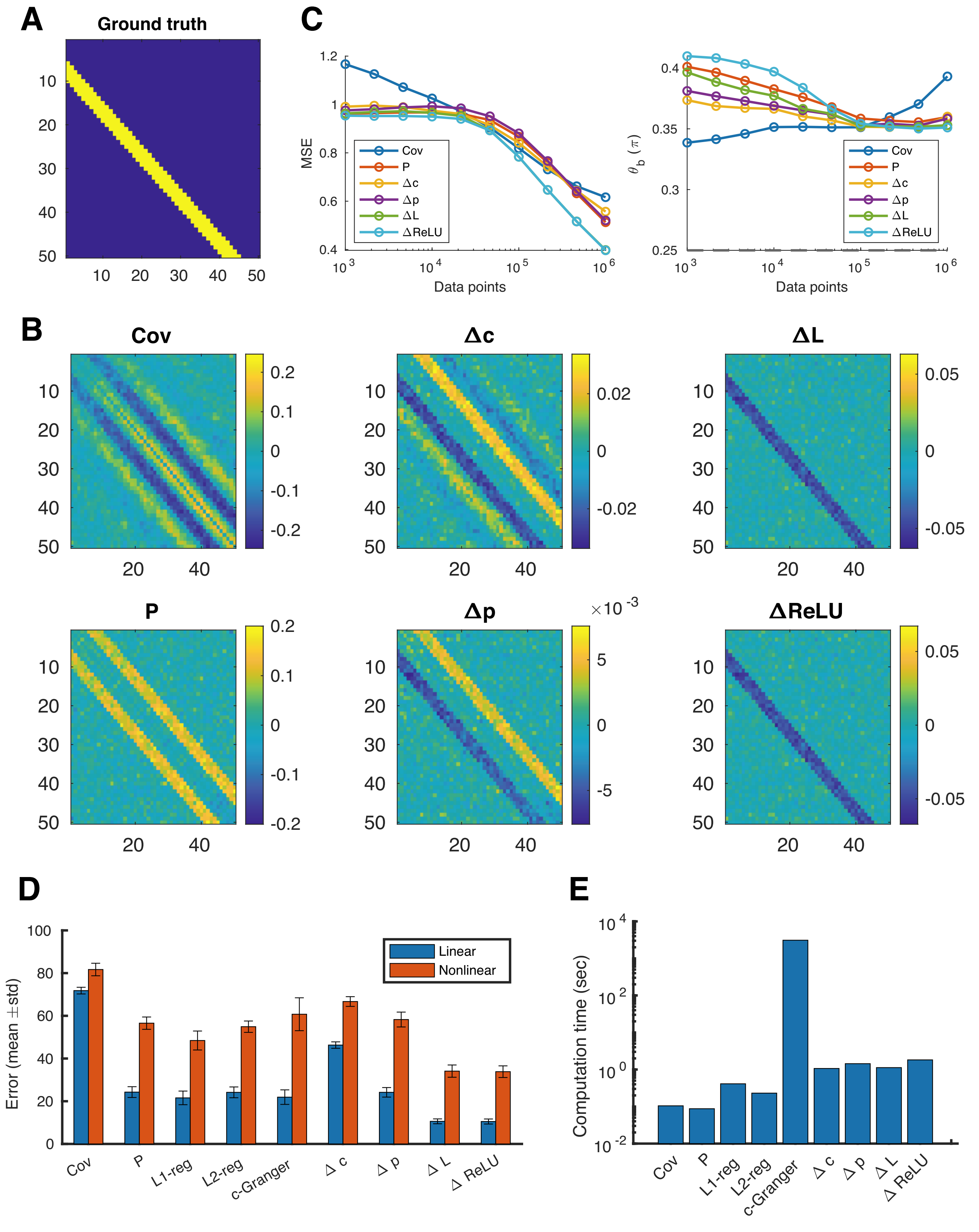


**Fig. S2. Estimator performance scaled up to larger networks.** (A) Ground truth connectivity pattern of a 50-node network. The connected edges were shown as yellow entries and they were assigned negative strength to stabilize simulation. (B) Estimated matrices using a sufficiently large data volume ($10^6$ data points) and nonlinear dynamics (Eq.24), which is the harder scenario according to Fig.S1. $\Delta$L and $\Delta$ReLU had cleaner estimation of the ground truth matrix. (C) Influence of dataset size on performance. (D) Estimation performance of both linear and nonlinear simulation. (E) Computation time in seconds.

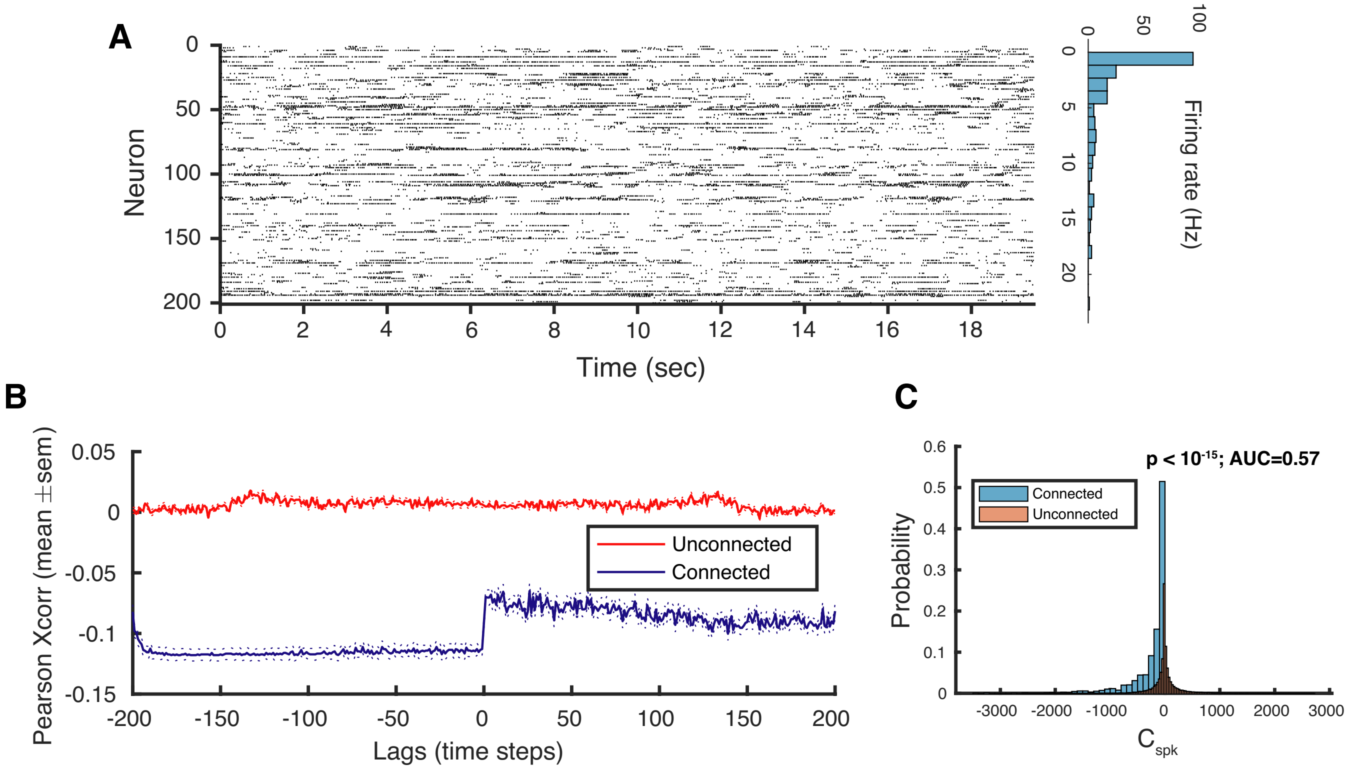


**Fig. S3. Connectivity estimation using spike trains** (A) Left: The spike train of 200 simulated neurons for 20 seconds (network sparsity: 0.04, connection strength: -0.4). Right: Histogram of the firing rate. (Average: 3.5Hz, Quiet neurons: 29). (B) Pearson cross correlogram of connected pairs and unconnected pairs. $C_{\mathrm{spk}}$ was calculated by averaging the cross correlation for the negative time lags. (C) Connected pairs have significantly (rank-sum $p < 10^{-15}$) smaller $C_{\mathrm{spk}}$ values compared to unconnected pairs. But the AUC (=0.57) value remained to be low.

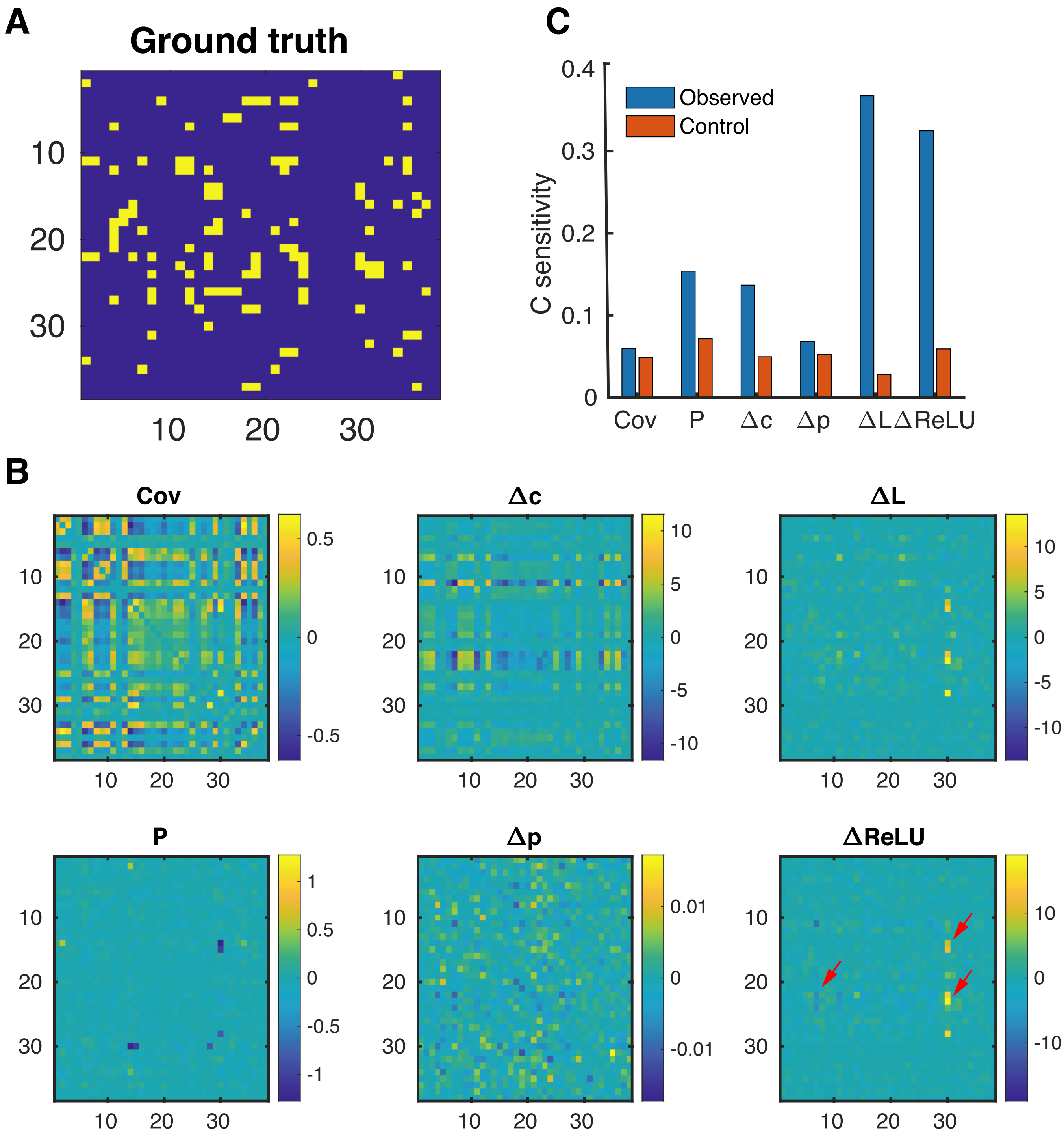


**Fig. S4. DDC is sensitive to strong anatomic connections at macroscopic level** (A) Intra-hemisphere strong connections (yellow entries) identified by Diffusion Spectrum Imaging (Node definition followed from Hagmann et al (1)). Time traces were generated using using both inter- and intra-hemisphere connections. (B) Estimated intra-hemisphere connections. Note the estimated strong connections (red arrow) in ΔL and ΔReLU correspond to true connections in (A). (C) Performance quantified by c-sensitivity. Control values were evaluated by comparing the estimation to a shuffled ground truth matrix.

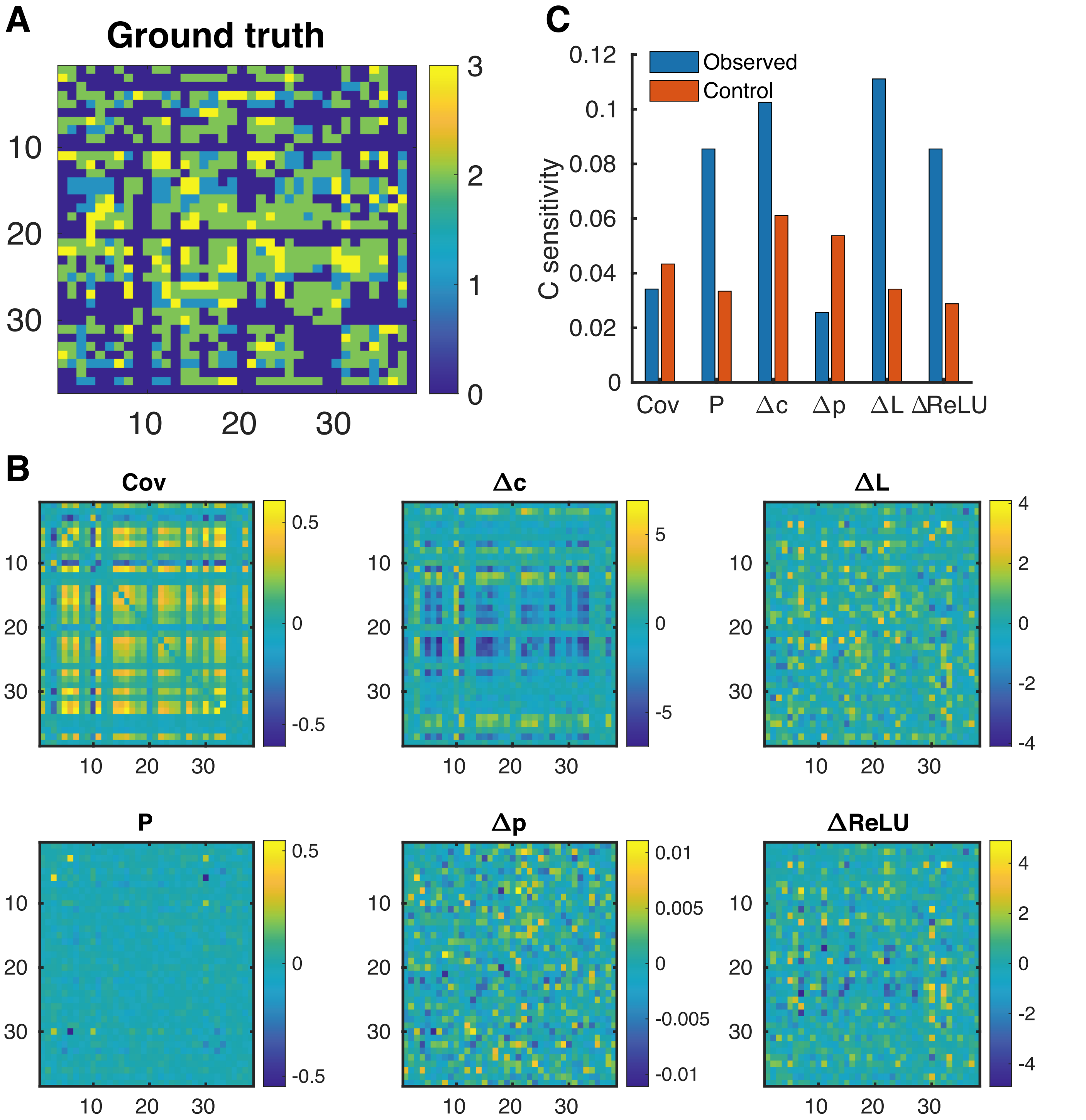


**Fig. S5. Macroscopic model supported by graded anatomical connections** (A) Intra-hemisphere anatomical connectivity. (B) Estimated intra-hemisphere connections. (C) Performance quantified by c-sensitivity. The overall low c-sensitivity indicates the difficulty of connection estimation.

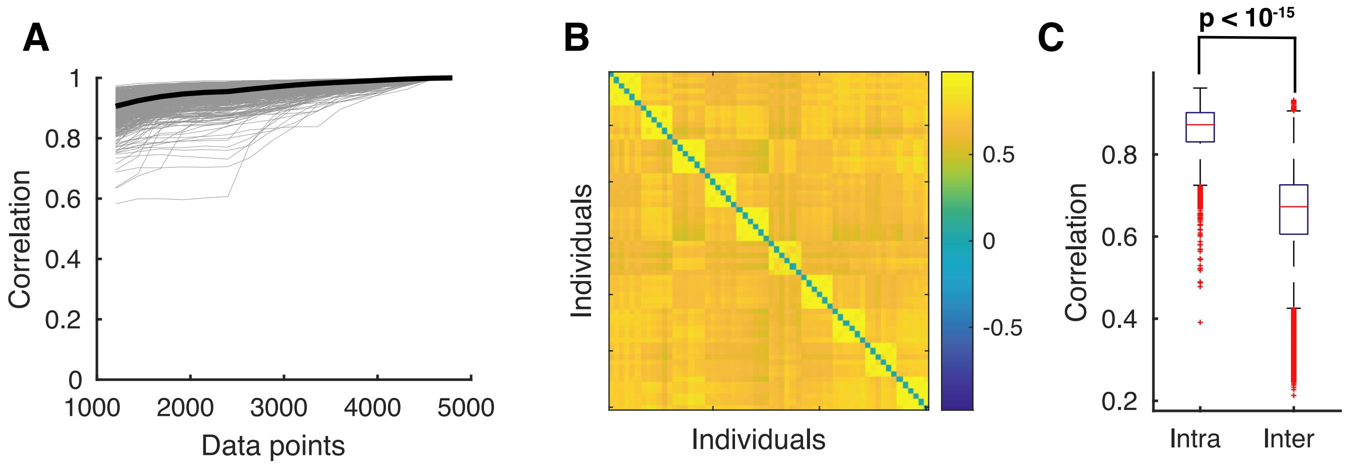


**Fig. S6. Intra- and inter-individual variability of sample covariance estimation across scan session.** A) Correlation of the full-length covariance estimation with covariance estimated using different amount of data. One scan session includes 1200 data points. Each gray line represents one subject and the black line is the average. B) Correlation of covariance estimated using a concatenation of two scan session within and between 10 randomly selected subjects. The block diagonal structure indicates a higher level of intra-individual similarity. C) Boxplot of intra- and inter-individual correlation pooled from all 1003 subjects (two-sided Wilcoxon rank sum test, p$< 10^{-15}$).

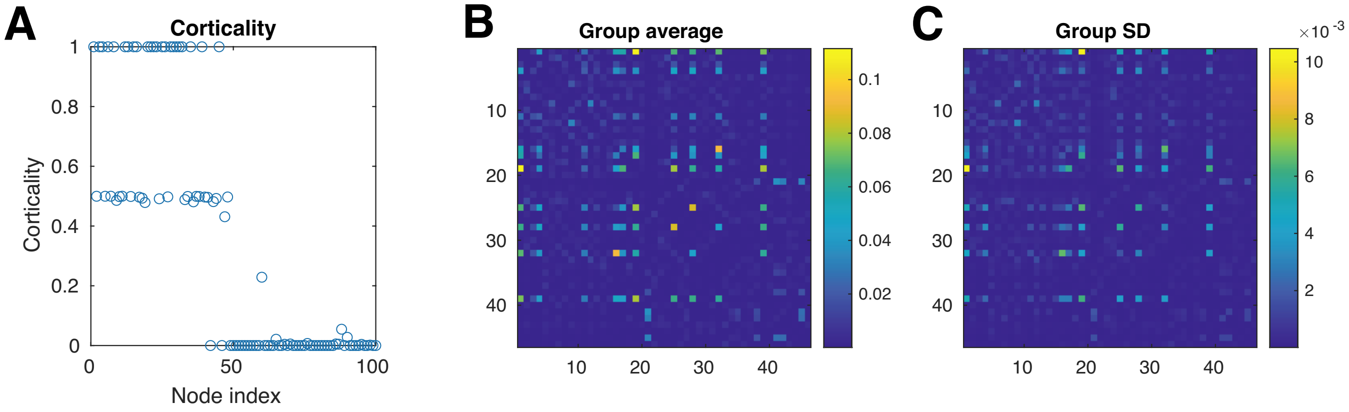


**Fig. S7. Individual level dMRI statistics.** (A) Corticality was defined as the proportion of cortical voxels within each IC. Since dMRI measurements are only available for cortical surface voxels. Our analysis was restricted to the first 46 ICs with corticality greater than 40%. (B) Average of the dMRI matrices across the entire 998 subjects. (C) Standard deviation.

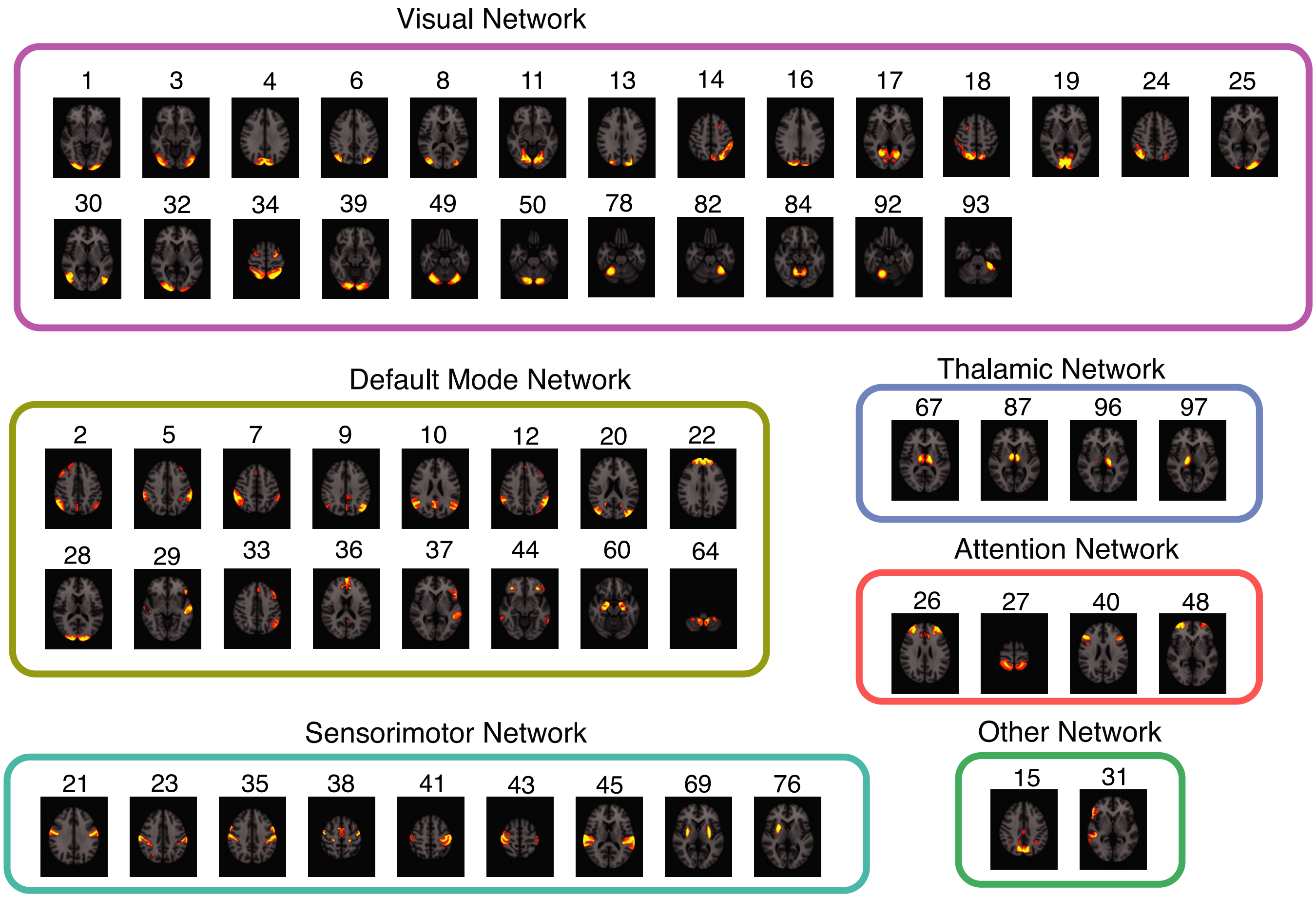


**Fig. S8. Group ICA parcellation shown on an MRI template**
